# IVT-seq reveals extreme bias in RNA-sequencing

**DOI:** 10.1101/005371

**Authors:** Nicholas F. Lahens, Ibrahim Halil Kavakli, Ray Zhang, Katharina Hayer, Michael B. Black, Hannah Dueck, Angel Pizarro, Junhyong Kim, Rafael Irizarry, Russell S. Thomas, Gregory R. Grant, John B. Hogenesch

## Abstract

**Background:** RNA sequencing (RNA-seq) is a powerful technique for identifying and quantifying transcription and splicing events, both known and novel. However, given its recent development and the proliferation of library construction methods, understanding the bias it introduces is incomplete but critical to realizing its value.

**Results:** Here we present a method, in vitro transcription sequencing (IVT-seq), for identifying and assessing the technical biases in RNA-seq library generation and sequencing at scale. We created a pool of > 1000 *in vitro* transcribed (IVT) RNAs from a full-length human cDNA library and sequenced them with poly-A and total RNA-seq, the most common protocols. Because each cDNA is full length and we show IVT is incredibly processive, each base in each transcript should be equivalently represented. However, with common RNA-seq applications and platforms, we find ∼50% of transcripts have > 2-fold and ∼10% have > 10-fold differences in within-transcript sequence coverage. Strikingly, we also find > 6% of transcripts have regions of high, unpredictable sequencing coverage, where the same transcript varies dramatically in coverage *between* samples, confounding accurate determination of their expression. To get at causal factors, we used a combination of experimental and computational approaches to show that rRNA depletion is responsible for the most significant variability in coverage and that several sequence determinants also strongly influence representation.

**Conclusions:** In sum, these results show the utility of IVT-seq in promoting better understanding of bias introduced by RNA-seq and suggest caution in its interpretation. Furthermore, we find that rRNA-depletion is responsible for substantial, unappreciated biases in coverage. Perhaps most importantly, these coverage biases introduced during library preparation suggest exon level expression analysis may be inadvisable.

## Background

High-throughput sequencing of RNA (RNA-seq) is a powerful suite of techniques to understand transcriptional regulation. Using RNA-seq, not only can we perform traditional differential gene expression analysis with better resolution, we can now comprehensively study alternative splicing, RNA editing, allele specific expression, and identify novel transcripts, both coding and non-coding RNAs [1–3]. In contrast to the more established microarray based RNA expression analysis, the flexibility of RNA-seq has allowed for the development of many different protocols aimed at different goals (e.g. gene expression of poly adenylated transcripts, small RNA sequencing, total RNA sequencing, etc.). However, this same flexibility has the potential for complex technical bias, as different methods are routinely employed in RNA isolation, size selection, fragmentation, conversion to cDNA, amplification, and finally, sequencing [4–7]. While progress has been made in generating and analyzing RNA-seq data, we understand comparatively little about the technical biases the various protocols introduce. Understanding these biases is critical to differential analysis, to avoiding experimental artifacts (e.g. in characterizing RNA editing), and to realizing the full potential of this powerful technology.

Previous efforts at understanding bias identified several contributing sources, including GC-content and PCR enrichment [8, 9], priming of reverse transcription by random hexamers [10], read errors introduced during the sequencing-by-synthesis reaction [11], and bias introduced by various methods of ribosomal RNA (rRNA) subtraction [7]. Studies that revealed these sources of bias typically use computational methods on existing sequencing data to assess the performance of various sequencing technologies and library protocols. One downside to this approach is that it can be difficult to know whether anomalies in coverage are natural, or are due to technical artifacts. For example, nearly every RNA-seq study has differences in intra-exonal coverage, which could arise from naturally occurring splice variants sharing part of an exon, or could be due to technical error in library construction or sequencing.

Given that researchers are continually developing new sequencing methodologies and library generation protocols [12], we need a means for assessing the technical biases introduced by each new iteration in technology. One attractive alternative is to generate libraries from RNA that has been *in vitro* transcribed (IVT) from cDNA clones, where the nucleotide sequence at every base is known, the splicing pattern established and inviolate, and the expression level is known to be uniform across the transcript. Thus, any observed biases in coverage or expression must be technical rather than biological. This is the experimental equivalent of simulated data that computational researchers commonly use to develop and assess alignment algorithms [13–15]. Jiang and colleagues used a similar approach with 96 synthetic sequences derived from *Bacillus subtilis* or the deep-sea vent microbe *Methanocaldococcus jannaschii* genomes [16], organisms that do not have RNA splicing or poly adenylation. The focus of that work, though, was creating a useful set of standards that could be used in downstream analysis, not exploring library construction bias in a comprehensive set of complex mammalian samples.

Here we present and apply IVT-seq at scale to better understand bias introduced by RNA-seq. In brief, individual plasmids were produced, pooled, and subjected to in vitro transcription. Next, this RNA was mixed with complex mouse total RNA at various concentrations, and sequenced using the two most common RNA-seq protocols, polyA seq or total RNA seq, on the Illumina platform. We find coverage bias in most IVT transcripts, with over 50% showing > 2-fold changes in within-transcript coverage and 10% having > 10 fold differences attributable to library preparation and sequencing. Additionally, we find > 6% of IVT transcripts contain regions of high, unpredictable sequencing coverage, which vary significantly between samples. These biases are highly reproducible between replicates and suggest that exon-level quantification may be inadvisable. Furthermore, we created sequencing libraries from the original plasmid templates and using several different RNA selection methods (rRNA depletion, polyA selection, and no selection). We find that both rRNA depletion and polyA selection are responsible for a significant portion of this coverage bias, and computational analysis shows that poorly represented regions of transcripts are associated with low complexity sequences. Taken together, these results show the utility of the IVT-seq method for characterizing and identifying the sources of coverage bias in sequencing technologies.

## Results and discussion

### IVT-seq library preparation and sequencing

To generate IVT-seq libraries (for full details, please see the *Materials and methods* section), individual glycerol stocks each harboring a single, human, fully sequenced plasmid from the Mammalian Gene Collection [17] were produced and plasmid DNA was extracted and plated at 50 ng per well in 384-well plates. The contents of three 384-well plates containing a total of 1062 cDNA clones (Additional file 10) were mixed, transformed into bacteria, and plated as single colonies. These plates were scraped, amplified for a few hours in liquid culture, and purified as a pool (Figure 1A). Next, plasmids were linearized, purified, and SP6 polymerase was used to drive *in vitro* transcription of the cloned cDNA sequences (Figure 1B). Following a DNase I treatment to remove the DNA template and RNA purification, a pool of 1062 different human RNAs derived from fully sequenced plasmids was produced.

**Figure 1:**
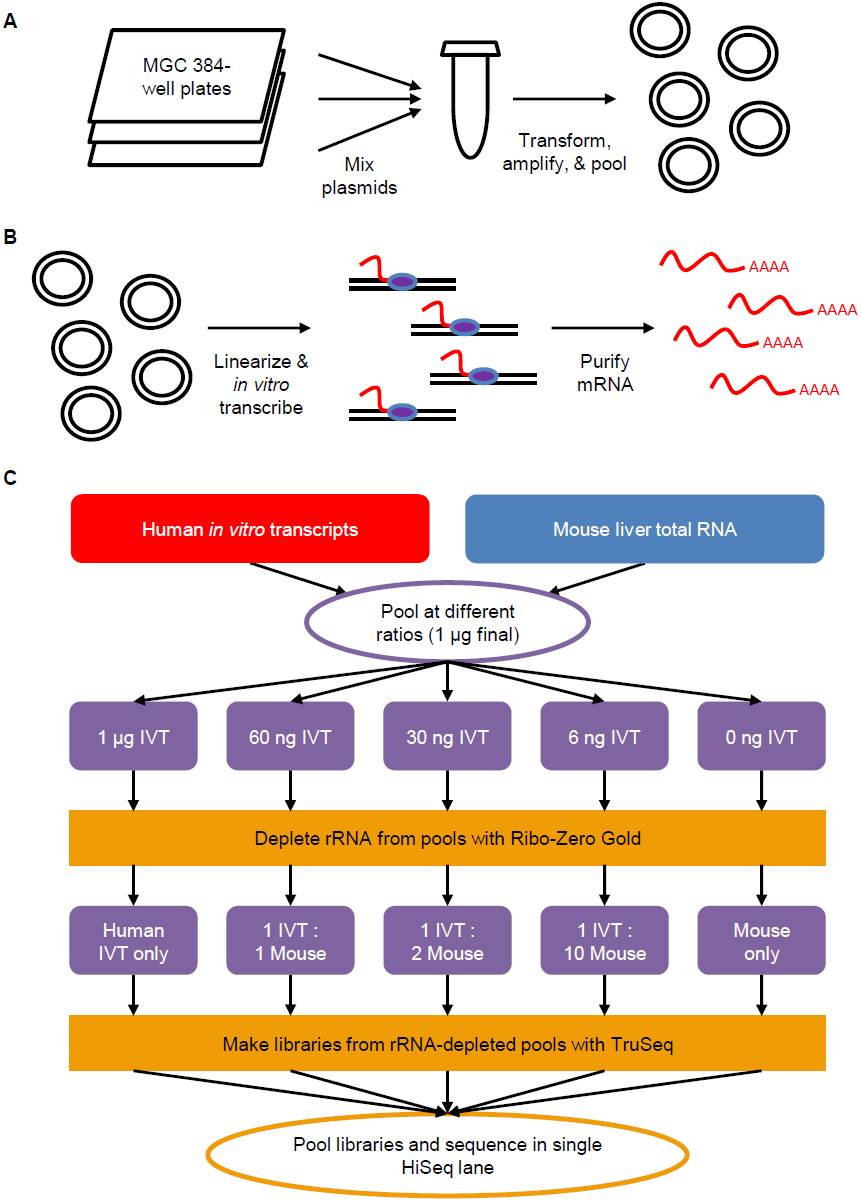
Construction of IVT-seq libraries. **(A)** Prepare pool of 1062 human cDNA plasmids. Contents of three 384-well plates containing MGC plasmids were pooled together. Pool was amplified via transformation in *E*. *coli*, and resulting clones were purified, and re-pooled. **(B)** Generate IVT transcripts. Pool of MGC plasmids were linearized and used a template for an *in vitro* transcription reaction. Enzymes and un-incorporated nucleotides were purified, leaving pool of poly(A) transcripts. **(C)** Create IVT-seq libraries. Listed quantities of IVT RNA were mixed with mouse liver total RNA to create six pools with final RNA quantities of 1 µg. Ribosomal RNA was depleted from these pools using the Ribo-Zero Gold kit. IVT RNA and mouse RNA are now present in pools at the listed ratios, following depletion of rRNA from mouse total RNA. These pools were used to generate RNA-seq libraries using Illumina’s TruSeq kit/protocol. This entire process was performed in duplicate. Replicate libraries were pooled separately and sequenced in separate HiSeq 2000 lanes (two lanes total).

To approximate what happens in a total RNA sequencing reaction, we subjected this IVT RNA to rRNA-depletion and then prepared libraries using the Illumina TruSeq protocol (Figure 1C, IVT only). To account for possible carrier effects, we also mixed the IVT RNA with various amounts of mouse total RNA derived from liver. The addition of the mouse RNA gave these samples greater diversity (transcripts from ∼10k genes vs. 1062) and more closely resembled a real biological sample. Also, by adding background RNA from a different species (mouse) than the IVT RNA (human), we make it easier to differentiate between the IVT transcripts and mouse sequences during downstream analysis. Since the IVT RNA does not contain rRNA sequences while the mouse RNA does, the quantity of mouse RNA will be significantly reduced by the rRNA depletion step. In order to account for this we mixed IVT and mouse RNA such that following rRNA depletion we would have final pools with IVT:mouse ratios of 1:1, 1:2, and 1:10. Finally, to account for mouse RNAs potentially mapping to the human reference genome and our IVT sequences, we prepared a pool consisting of mouse RNA alone. We pooled the resulting six libraries and sequenced them using an Illumina HiSeq 2000. We performed this entire process in duplicate.

### Mapping and coverage of IVT-seq data

Following sequencing and de-multiplexing, we aligned all of the data to the human reference genome (hg19) using the RNA-seq Unified Mapper (RUM) [14]. For all analyses, we only used data from reads uniquely mapped to the reference, excluding all multi-mappers (data contained in RUM_Unique and RUM_Unique.cov files). Of the 1062 original IVT transcripts, we found 11 aligned to multiple genomic loci, while 88 aligned to overlapping loci. To avoid any confounding effects in our analyses, we filtered those transcripts from all analyses, leaving us with 963, non-overlapping, uniquely-aligned IVT transcripts. We saw excellent correlation in expression levels between replicates (transcript-level R^2^ between replicates > 0.95; Additional file 1: Figure S1A). Secondly, at least 90% of the 963 IVT transcripts are expressed with an FPKM ≥ 5 in all IVT-seq datasets except mouse only (Table 1). In the IVT-only samples, over 80% of the IVT sequences are expressed above 100 FPKM (Additional file 1: Figure S1B). Since we prepared the MGC plasmids and IVT transcripts as pools, it is likely that the IVT transcripts showing low or zero coverage were initially present at low plasmid concentrations prior to the transformation and IVT steps. Using the IVT-seq technique, we are able to specifically detect the vast majority of the human IVT transcripts with high coverage in both the absence and presence of the background mouse RNA.

**Table 1:** Detection of source cDNA sequences in IVT-seq.

|  |  |  |  |
| --- | --- | --- | --- |
| Total number of cDNA clones: | 963 |  |  |
|  | Replicate 1 | Replicate 2 |  |
| # of clones detected (FPKM $\geq 5$): | | | |
| IVT Only | 869 | 870 |  |
| 1:1 Mix | 877 | 876 |  |
| 1:2 Mix | 886 | 883 |  |
| 1:10 Mix | 896 | 892 |  |
| Mouse Only | 278 | 271 |  |
| PolyA Selection | 829 | - |  |
| No Selection | 870 | - |  |
| Plasmid Library | 924 | - |  |
| Average, normalized* depth of coverage for detected clones: |  |  |  |
| IVT Only | 76.09 | 80.22 |  |
| 1:1 Mix | 75.15 | 75.06 |  |
| 1:2 Mix | 65.79 | 69.40 |  |
| 1:10 Mix | 37.50 | 47.46 |  |
| Mouse Only | 01.58 | 02.42 |  |
| PolyA Selection | 72.27 | - |  |
| No Selection | 72.74 | - |  |
| Plasmid Library | 42.08 | - |  |

\*Average depth of coverage is normalized by the number of millions of fragments mapped to the human reference in each sample.

While we do see reads aligned to the human IVT transcripts in the mouse only data, these transcripts collectively represent ∼2% of reads (Table 1). Those transcripts with higher coverage are likely the result of mouse reads aligning to highly similar human sequences. We excluded these sequences from our analyses.

### Within-transcript variation in RNA-seq coverage of IVT transcripts

Consider first the IVT only data. Given that these transcripts were generated from an IVT reaction using cDNA sequences, this data is unaffected by splicing or other post-transcriptional regulation. Thus, most regions of transcripts should be “expressed” and present at similar levels. The exceptions would be repetitive sequences that map to multiple genome locations and may be poorly represented, and the ends of the cDNAs, which are subject to fragmentation bias. To account for this we created a simulated dataset which models the fragmentation process and which deviates from uniform data only by the randomness incurred by fragmentation. We generated two such datasets using the Benchmarker for Evaluating the Effectiveness of RNA-Seq Software (BEERS) [14]. The first dataset contains all of the IVT transcripts expressed at roughly the same level of expression (∼500 FPKM). For the second, we used FPKM values from the IVT-only samples as a seed, creating a simulated dataset with expression levels matching real data (Additional file 2: Figure S2). These datasets are referred to as simulated and QM-simulated (Quantity Matched), respectively. The simulated data provides an ideal result, while the QM data allows us to control for any artifacts arising from expression level (eg. transcripts with lower expression may show more variability). Next, we aligned both simulated datasets using RUM, with the same parameters as for the biological data. Thus, both simulated datasets also serve as a controls for any artifacts introduced by the alignment (eg. low coverage in repeat regions). For full details on the creation of simulated data, see the *Materials and methods* section.

Using IVT data derived from the BC015891 transcript as a representative example, the ideal, theoretical coverage plot from the simulated data shows near-uniform coverage across the transcript’s entire length, with none of the extreme peaks and valleys characteristic of biological datasets (Figure 2A). However, our observed data shows a high degree of variability, with peaks and valleys within an exon (Figure 2B). Furthermore, these patterns are reproducible across our replicates (Additional file 3: Figure S3). We see many other cases of extreme changes in coverage; over 50% of the IVT transcripts show > 2-fold changes in within-transcript coverage attributable to library preparation and sequencing (Table 2 and Additional file 4: Figure S4). For example, BC009037 shows sudden dips to extremely low levels of expression in both of its exons (Figure 2C). Both simulated datasets show no such patterns, which indicates this coverage variability is not the result of alignment artifacts. Furthermore, the absence of this pattern in the QM-simulated data indicates these fold-change differences in coverage are not due to sampling noise introduced by transcripts with low or high coverage. In the case of BC016283, the peaks and valleys in coverage lead to greater than five-fold differences in expression levels between exons (Figure 2D). Once again, these patterns are reproducible across replicates (Additional file 3: Figure S3). The SP6 polymerase cannot fall off and then re-attach at a later point in the transcript, leaving a region un-transcribed. Therefore, given that these patterns show troughs followed by peaks, they cannot be the result of artifacts from *in vitro* transcription. Furthermore, we sequenced the IVT products directly and found the vast majority were transcribed with little to no bias. Taken together, these data suggest that these coverage patterns are primarily the result of technical biases introduced during library construction, rather than biology. These results are consistent with a previous study that uses *in vitro* transcribed RNA as standards in RNA-seq experiments [16], suggesting that our IVT-seq methodology is suitable for identifying technical variability in sequencing data.

**Figure 2:**
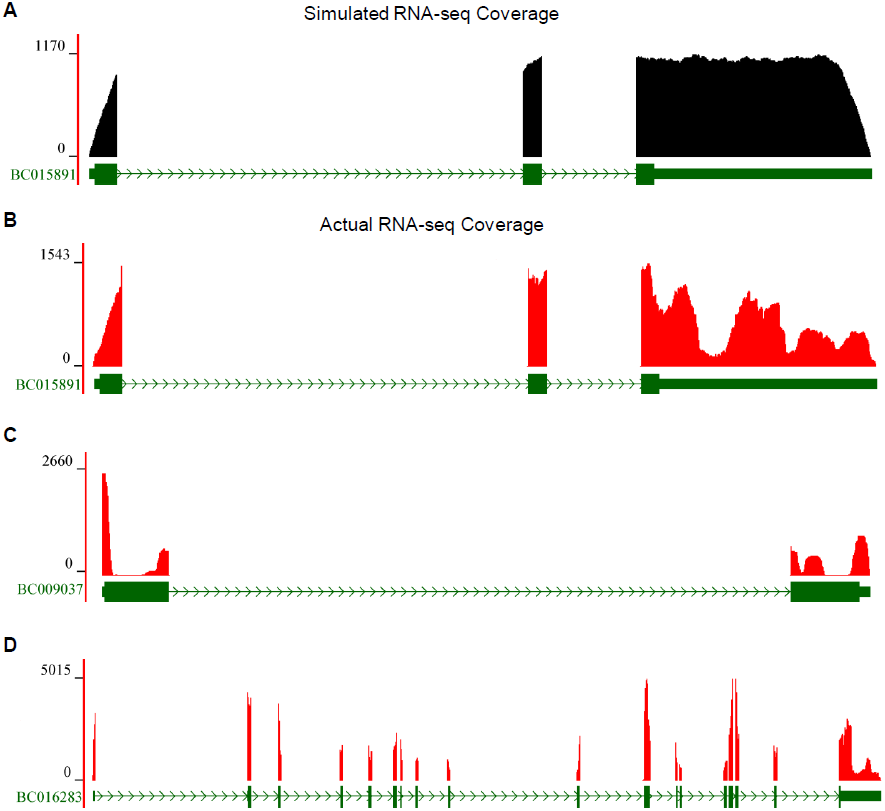
Within-transcript and between-sample variations in RNA-seq coverage. **(A)** Simulated RNA-seq coverage for a representative IVT transcript (BC015891). RNA-seq coverage plot (black) is displayed according to the gene model (green), as it is mapped to the reference genome. Blocks correspond to exons and lines indicate introns. The chevrons within the intronic lines indicated the direction of transcription. Numbers on y-axis refer to RNA-seq read-depth at a given nucleotide position. **(B)** The actual RNA-seq coverage plot for BC015891 in the IVT-only sample. Representative coverage plots for the IVT transcripts **(C)** BC009037 and **(D)** BC016283 are displayed according to the same conventions used above. All transcripts are displayed in the 5′ to 3′ direction.

**Table 2:** Fold-change differences in within-transcript coverage by library type.

|  | # of IVT-transcripts with fold-change differences: |  |  |
| --- | --- | --- | --- |
|  | > 2 | > 10 | > 100 |
| rRNA-Depleted | 713 (74.0%) | 110 (11.4%) | 17 (1.7%) |
| PolyA Selection | 678 (70.4%) | 163 (16.9%) | 7 (0.7%) |
| No Selection | 400 (41.5%) | 31 (3.2%) | 3 (0.3%) |
| Plasmid | 189 (19.6%) | 14 (1.5%) | 3 (0.3%) |
| Simulated | 0 | 0 | 0 |
| QM-Simulated | 0 | 0 | 0 |

The plasmid data provides a measure of bias from library preparation/sequencing, while the ‘no selection’ data accounts for potential artifacts from the *in vitro* transcription step. To calculate the percentage of transcripts affected by bias due specifically to library preparation and sequencing, but not sequence or *in vitro* transcription artifacts, we perform the following calculation: rRNA-depletion % - no selection % + plasmid %. So we find 74% – 41.5% + 19.6% = 52.1% of transcripts in the rRNA-depleted data have > 2-fold difference in coverage, and 11.4% – 3.2% + 1.5% = 9.7% have > 10-fold difference in coverage.

### Between-sample variation in RNA-seq coverage of IVT transcripts

In addition to this variability within transcripts, we also find many transcript regions showing extreme variability in coverage across samples (Figure 3). For example, the sixth exon of BC003355 varies wildly relative to the remainder of the transcript across all IVT:mouse dilutions. Interestingly, the overall pattern of variation relative to the rest of the transcript across the dilutions is maintained between the replicates. Almost no reads in the mouse-only sample map to this transcript, which eliminates the possibility that this variability is due to incorrect alignment of mouse RNA.

**Figure 3:**
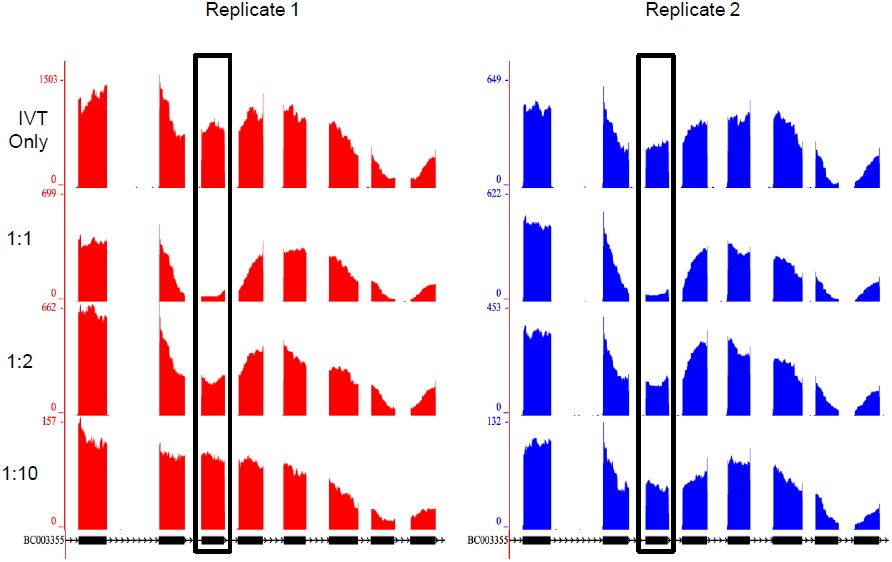
Between-sample variations in RNA-seq coverage. RNA-seq coverage plots across all samples for exons 4 – 11 of the IVT transcript BC003355. The black rectangles identify exon six, which shows extreme variability in coverage relative to the rest of the transcript when viewed across all of the samples. The ratio of IVT RNA to moue RNA is listed to the left of each sample’s coverage plots. Coverage plots (red for first replicate; blue for second replicate) are displayed according to the gene model (black), as it is mapped to the reference genome. Blocks in the gene model correspond to exons and lines indicate introns. The chevrons within the intronic lines indicated the direction of transcription. Numbers on y-axes refer to RNA-seq read-depth at a given nucleotide position.

Including BC003355, we find 86 regions of high, unpredictable coverage (hunc) spread across 65 transcripts (Additional file 11). Therefore, over 6% of the 963 IVT transcripts contain regions showing wild but reproducible variations in RNA-seq coverage between samples. While identifying these hunc regions, we used a two-stage filter to eliminate variable regions resulting from mouse reads mapped to highly similar human sequences. First, we eliminated all hunc regions coming from transcripts with FPKM >= 5 in either mouse-only dataset. Next, to account for localized misalignment of mouse reads, we filtered out all hunc regions with an average coverage >= 10 in either mouse-only dataset. We also removed those hunc regions with mouse-only coverage >= 10 in the flanking 100bp on either side. Given the stringent criteria we used to identify these hunc regions (see *Materials and methods* section for full details), it is likely that this is an underestimate. To address the possibility that mouse RNAs may interact with homologous human RNAs and interfere with them in *trans*, we assayed the sequences surrounding these regions using the MEME Suite [18], but we found no sequence motifs these regions have in common. Furthermore, the depth of coverage at these regions does not follow a linear relationship with the increasing mouse RNA, which suggests it is not simply a direct interaction with the background RNA. There is no clear cause for these hunc regions, particularly since we prepared all samples from the same pool of IVT RNA and the only difference between samples is the relative ratios of IVT RNA to mouse liver RNA. We also searched for hunc regions that were divergent between the two replicates, but found none. If such regions do exist, they could be identified and overcome by creating libraries in duplicate. The hunc regions we identified above with expression patterns maintained between replicates present a greater challenge, as they could not be identified and filtered out by creating duplicate libraries. This is particularly problematic for using exon-level expression values to identify alternative splicing events or differential expression. The within-transcript and between-sample variation we see in our IVT-seq data suggests that library generation introduces strong technical biases, which could confound attempts to study the underlying biology.

### Sources of variability in RNA-seq coverage

There are three potential sources for technical bias in library preparation: RNA-specific molecular biology (i.e. RNA fragmentation, reverse-transcription), RNA selection method (i.e. rRNA-depletion, polyA selection), and sequencing-specific molecular biology (i.e. adapter ligation, library enrichment, bridge PCR). To identify biases introduced solely by sequencing-specific molecular biology, we created a DNA-seq library from the same MGC plasmids used as templates for the IVT-seq libraries (Additional file 5: Figure S5). In doing this, we skip the steps specific to the IVT or RNA molecular biology. We also prepared two additional IVT-seq libraries using polyA selection or no selection, instead of rRNA depletion. By comparing our plasmid library data and the IVT-seq data using various selection methods, we can identify which coverage patterns are the result of RNA-specific molecular biology, the RNA selection method, or of some common aspect of the library generation protocol.

We sequenced the plasmid library using an Illumina MiSeq and aligned the resulting data to the human reference genome using the same method as the IVT-seq libraries. In this plasmid data, we see 924 of the cDNA clone sequences with FPKM values ≥ 5, compared to ∼870 in both of the IVT only samples (Table 1). This small drop in coverage is likely because the IVT RNA goes through more pooling steps during library construction than the plasmids. Furthermore, the plasmids are not affected by transcription and reverse transcription efficiencies. Additionally, the plasmid data maps to the cDNA sequences with an average, normalized coverage of 42.08, which is within the range of coverage values we see for the IVT-seq samples. We sequenced the no selection and polyA selection libraries on a HiSeq 2500. This data also shows cDNA clone coverage values similar to the other IVT-seq libraries.

The plasmid data represents the “input” into the IVT reaction and the no selection data represents the closest measure of its direct output. By measuring the 3’/5’ ratio in depth of coverage for each IVT transcript, we can assess the processivity of the SP6 polymerase. In a perfectly processive reaction, this 3’/5’ ratio would be 1, indicating the polymerase did not fall off the cDNA template and lead to the formation of truncated products. The median 3’/5’ ratios for the plasmid and no selection data were 1 and 0.98, respectively, indicating premature termination of the IVT reaction was not a factor in our analyses.

### Effect of different RNA selection methods on coverage patterns

Our analysis is illustrated by an examination of the coverage plots for BC003355 across all of our different datasets. The high degree of variability we noted in this gene’s coverage plot from our rRNA-depleted data is absent in the no selection and plasmid data (Figure 4A). While the polyA data also shows fewer peaks and valleys than the rRNA depleted total RNA-seq data, it is marked by the well-documented 3’ bias. This data suggests that the rRNA-depletion step is likely responsible for a large quantity of the observed coverage biases.

**Figure 4:**
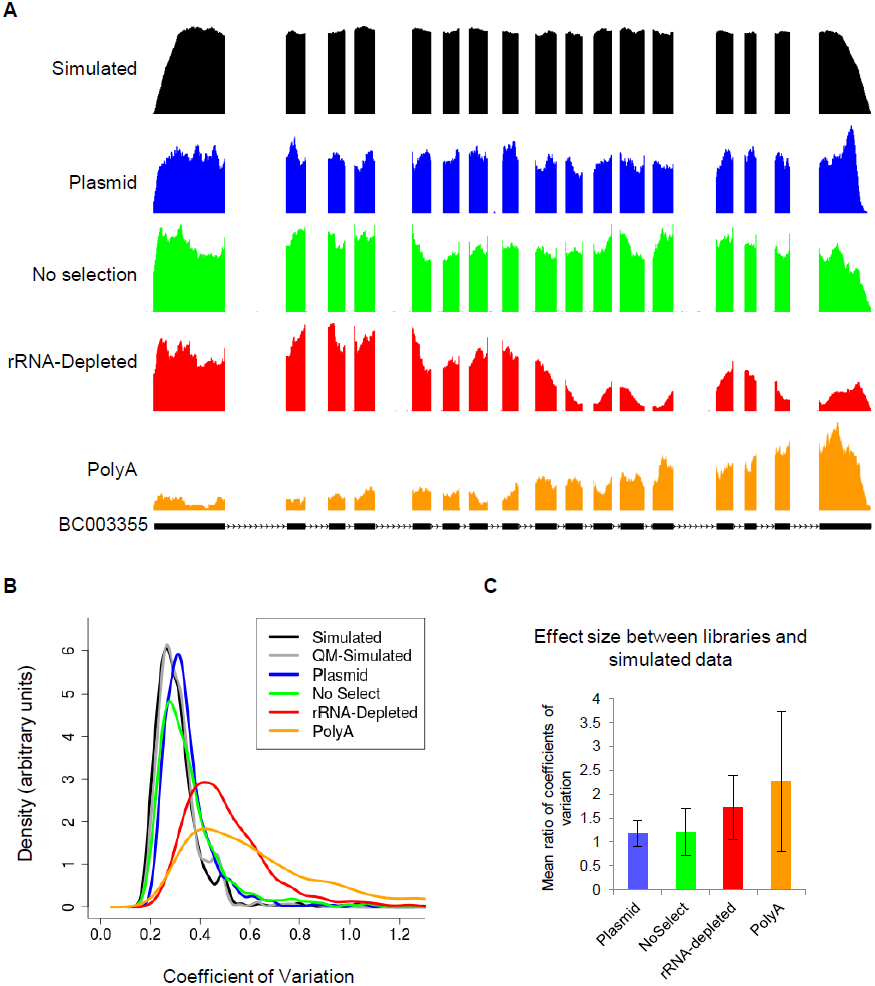
Sources of bias in RNA-seq coverage. **(A)** RNA-seq coverage plots for IVT transcript BC003355 from simulated (black), plasmid (blue), no selection (green), rRNA-depleted (red), and polyA (orange) data. The gene model is displayed in black, below all of the coverage plots. Blocks correspond to exons and lines indicate introns. The chevrons within the intronic lines indicated the direction of transcription. **(B)** Distributions for coefficients of variation across data displayed above, with the addition of the QM-simulated data (gray). Note that while the graph is cutoff at a coefficient of variation of 1.3, the tails for the Ribo-Zero and PolyA distributions extend out to 2.13 and 2.7, respectively. **(C)** Effect sizes for the differences in distribution of coefficients of variation between sequencing libraries and simulated data. Effect size is calculated as the mean of the per-transcript ratios of coefficients of variation between a given library and the simulated dataset. Error bars are the standard deviation of this ratio.

To quantify the variability for each selection method, we calculated the coefficient of variation at the single base level in coverage for all IVT transcripts across each of these datasets (Figure 4B). Using a Wilcoxon rank-sum test (plasmid n = 924, no selection n = 870, rRNA-depleted n = 869), we find the rRNA-depleted data has significantly higher variability than the no selection and plasmid data (*p* < 2.2e-16). Furthermore, the rRNA-depleted and polyA libraries are > 60% more variable on average than the plasmid library (Figure 4C). This suggests that a significant portion of the observed variability in coverage across transcripts in the IVT-seq data is the result of RNA-specific molecular biology, specifically the RNA selection step. Furthermore, after accounting for bias introduced by the sequences themselves (plasmid data) and bias introduced by the IVT reaction (‘no selection’ data), we find that 50% of transcripts have 2-fold and 10% have 10-fold variation in within transcript expression (Table 2 and Additional file 4: Figure S4). While it is well appreciated that polyA selection introduces bias, we found that rRNA-depleted data introduced just as much if not more. Neither simulated dataset showed transcripts with a 2-fold or higher change in within transcript expression. Again, this suggests that the observed within transcript variations are not the result of alignment artifacts or sampling due to low/high expression. One commonly acknowledged source of bias arises from random priming during library preparation [10]. When we examined the different libraries, we saw that fragments from all of the RNA-seq data showed nucleotide frequencies characteristic of random priming bias (Additional file 6: Figure S6). As expected, the plasmid data showed no such bias, since it was derived directly from DNA and required cDNA generation step. However, the significant differences between all RNA libraries suggest that bias from random priming is not the only factor. The plasmid and no selection data still contain a fair amount of variability when viewed alongside the simulated data (Figure 4A; black). When we examine the entire dataset, both the plasmid and no selection data have significantly higher variation than either simulated dataset (Wilcoxon rank-sum test; simulated data n = 963, QM-simulated data n = 869, plasmid n = 924, no selection n = 870; *p* < 2.2e-16). This data suggests that sequencing-specific molecular biology common to all libraries we prepared (adapter ligation, library amplification via PCR), is also responsible for a portion of the observed coverage variability and sequencing bias.

### Biases associated with sequence features are dependent on RNA selection method

Given these significant differences in coverage variability, we sought to identify sequence features that might contribute to this bias. We considered three quantifiable sequence characteristics: hexamer entropy, GC-content, and sequence similarity to rRNA (see *Materials and methods* for a detailed description of these metrics). For each sequencing strategy (plasmid, no selection, rRNA-depleted, polyA), we tested if any of the three sequence characteristics has a significant effect on variability in sequencing coverage, as measured by the coefficient of variation. While we are primarily focused on coverage variability as an indicator of sequencing bias, we also looked at depth of coverage, as measured by FPKM.

For each sequencing strategy we sorted the transcripts by coverage variability or depth. Next, we selected the 100 most and 100 least extreme transcripts from each list. We compared the values of the sequence characteristics between the 100 most and 100 least extreme transcripts using a Wilcoxon rank-sum test. Significant *p*-values indicate a significant association of the sequence characteristic with coverage variability and/or depth. The results of our analysis are displayed as box-plots (Figure 5 and Additional file 8: Figure S8).

**Figure 5:**
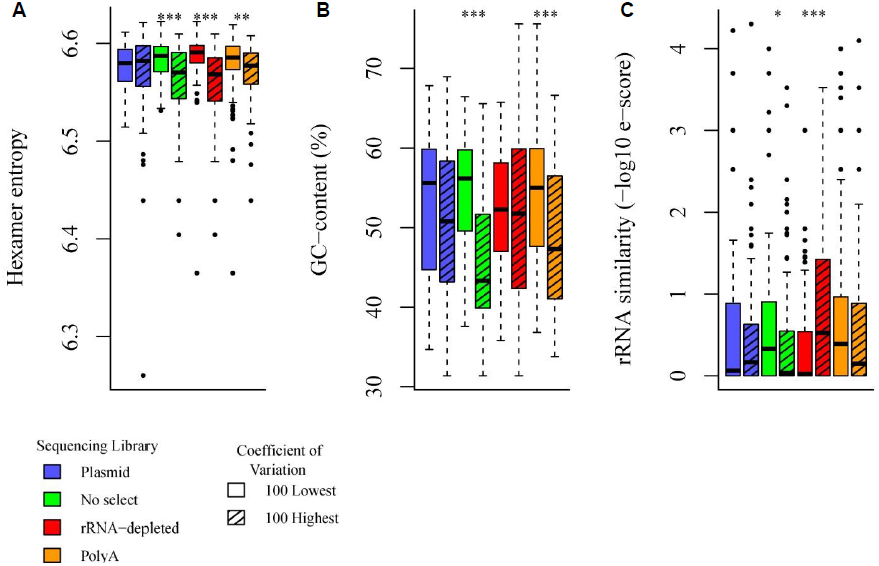
Effects of sequence characteristics on coverage variability. Distributions of **(A)** hexamer entropy, **(B)** GC-content, and **(C)** rRNA sequence similarity for the 100 transcripts with the highest and lowest coefficients of variation for transcript coverage from the plasmid, no selection, rRNA-depleted, and polyA libraries. Asterisks indicate the significance of a Wilcoxon signed-rank test comparing values for the listed sequence characteristics between each pair of groups from the same sample. * = *p*-value < 0.05; ** = *p*-value < 0.01; *** = *p*-value < 0.001.

To check for any confounding effects between coverage depth and variability, we tested the least and most expressed transcripts for any correlations with variability in coverage (Additional file 7: Figure S7). The polyA library showed a significant correlation (*p* < 2.2e-16) between coverage variability and depth, which indicates sequence features could be affecting coverage through variability (or vice versa). The rRNA-depleted data showed a slight, significant correlation (*p* = 0.04933). It is possible some feature of RNA selection affects both variability and coverage, given that we saw no significant correlations for the two remaining samples.. This indicates that coverage variability and depth are independent for the plasmid and no selection data.

All three sequence characteristics have a significant association with variability and depth-of-coverage in at least one of the sequencing strategies. In particular, lower hexamer entropy, a measure of sequence complexity [19–21], is strongly associated with higher variance in all of the RNA libraries (no selection *p* = 4.712e-05; rRNA-depletion *p* = 3.956e-11; polyA *p* = 0.003921; Figure 5A). This suggests that bias associated with hexamer entropy is due partially to RNA-specific procedures in library preparation. Furthermore, an association with lower hexamer entropy indicates there are more repeat sequences in the transcripts with higher variability. This could be indicative of complex RNA secondary structures, as repeated motifs could facilitate hairpin formation. Furthermore, the absence of this association from the plasmid data suggests that this observation is not due to mapping artifacts. The plasmid data contains the same sequences as the RNA-seq data, and would be subject to the same biases introduced by our exclusion of multi-mapped reads.

Higher GC-content is strongly associated with lower coverage variability in the no selection and polyA data (*p* = 5.627e-13; *p* = 4.914e-05; Figure 5B), suggesting that the effects of GC-bias on within-transcript variability could arise, in part due to some RNA-specific aspects of library preparation. Also, it appears that GC-bias is not a significant contributing factor to either depth of coverage, or the extreme variability in the rRNA-depleted data. Meanwhile, lower GC-content is associated with higher coverage in the plasmid data (*p* = 3.776e-05), and lower coverage depth in the no selection and polyA libraries (no selection *p* = 8.531e-05; polyA *p* = 0.0009675; Additional file 8: Figure S8B). Given that this trend switches directions between the plasmid library and the RNA libraries, this also suggests that some RNA-specific aspect of library preparation is introducing GC-bias distinct from the high GC-bias associated with Illumina sequencing [22].

Interestingly, higher rRNA sequence similarity is associated with higher coverage variability in the rRNA-depleted library (*p* = 9.006e-05) and lower variability in the no selection library (*p* = 0.0367; Figure 5C). It is unsurprising that similarity to rRNA sequences contributes to variability in the rRNA-depleted data, given that rRNA-depletion is based upon pair-binding between probes and rRNA sequences. While it is unclear why this trend is reversed in the no selection library, it is striking given the significant increase in within-transcript variability between the no selection and rRNA-depleted libraries (Figure 4B). Furthermore, we see a slight but highly significant correlation (Pearson R^2^ = 0.308452; *p* < 2.2e-16) between a transcript sequence’s similarity to rRNA, and the magnitude of the difference in coverage between the no selection and rRNA-depleted libraries (Additional file 9: Figure S9). While the majority of the factors contributing to the extreme bias in sequence coverage we see in the rRNA-depleted data remain unclear, our data suggests this is could be partially due to depletion of sequences homologous to rRNA.

Taken together, all of our data demonstrates the utility and potential of the IVT-seq method to identify sources of technical bias introduced by sequencing platforms and library preparation protocols.

## Conclusions

In this study, we present IVT-seq as a method for assessing the technical variability of RNA sequencing technologies and platforms. We created a pool of *in vitro* transcribed RNAs from a collection of full length human cDNAs, followed by high-throughput sequencing (Figure 1). Since we know the identities and sequences of these IVT transcripts, and since they were created under conditions not affected by splicing and post-transcriptional modification, they are ideal for identifying technical biases introduced during RNA-seq library generation and sequencing. We used this method to demonstrate that library generation introduces significant biases in RNA-seq data, adding extreme variability to coverage and read-depth along the length of sequenced transcripts (Figure 2). Our most striking finding is that over 50% of the IVT transcripts show > 2-fold differences in this within-transcript coverage attributable to library preparation and sequencing, in the polyA and rRNA-depleted data (Table 2). We prepared all RNA-seq libraries from the same pool of IVT RNA, so these differences are due to library construction and sequencing methods. Furthermore, 6% of the IVT transcripts contain regions with high unpredictable coverage variability (huncs) across different dilutions of IVT and mouse liver RNA (Figure 3). We found it particularly concerning that these huncs are consistent between replicates, as this means these regions cannot be indentified and avoided by making replicate libraries. While the exact cause of this effect is unclear, it could be due to some trans interaction between different RNA (human IVT RNA and the background mouse liver RNA). If this is the case, it could prove difficult to account for, given the challenges we have already encountered making predictions for miRNA targets and RNA secondary structure. Based on these results, we strongly recommend caution in interpreting exon-level quantification data, particularly for identifying and quantifying alternative splicing events, without further understanding of these biases.

Using simulated data and by sequencing at various stages of the process (plasmids, unselected IVT RNAs, rRNA-depleted, and polyA selected), we found each step introduces bias. Regions of certain IVT transcripts are underrepresented in both DNA and RNA, suggesting something inherent in their structure may resist cloning and sequencing properly. The IVT reaction has its own biases, however, by and large, it worked extremely efficiently with 90%of the input templates producing transcripts at detectable levels. PolyA sequencing revealed the well described 3’ bias. Finally, we saw extreme bias introduced by the rRNA-depletion step. Though we have yet to find the majority of the sources for this extreme bias, knowing that it occurs and that is at least partially due to rRNA sequence similarity is an important first step. By making this data available to the community, we hope that new experimental and analysis methods can be developed to account for the biases inherent in various aspects of RNA-seq.

Moreover, IVT-seq could be more broadly employed. By itself, the MGC collection has cDNAs derived from more than 16,000 mouse and human genes, including hundreds of genes for which there are more than one form. Therefore, in principle, it is possible to generate sequence profiles for representatives for nearly 2/3 of the mammalian transcriptome, or spike in datasets to develop new and better methods for splice form detection and quantification. Similar profiling approaches could do the same for other organisms. In addition, IVT-seq is also immediately relevant to RNA-seq method development, e.g. developing new protocols or refining existing ones. Finally, the method is not specific to Illumina sequencing and could be used to account for bias in other sequencing chemistries and methods (e.g. SOLID, Ion Torrent, PacBio, etc.).

Importantly, we are not suggesting that current generation RNA-seq is not a fantastic new technology or that quantification data is incorrect, particularly given the validated, reproducible results researchers have been able to gain through its use. Rather, we wish to provide a cautionary note that our understanding of this technology is still relatively new and incomplete. It is our hope that through the use of this data and IVT-seq, we will develop the means to minimize or account for bias in RNA-seq and truly realize the vision of digital gene expression.

## Materials and methods

### Amplification of plasmid library

Glycerol stocks containing individual cDNAs (cloned into pCMV-Sport 6 plasmid) from the Mammalian Gene Collection [17], were produced and plasmid DNA was extracted and plated at 50 ng per well in 384-well plates. The contents of three 384-well plates (total of human 1062 transcripts; Additional file 10) were collected as follows: 10 µl sterile dH_2_O was added to each well and incubated at 37°C for 10 min to resuspend plasmid DNA in water. Plasmid DNAs were collected and combined in 1.5 mL tube with aid of multichannel pipette and concentrated by ethanol precipitation. To amplify the library 10 ng of plasmid library was transferred into *E.coli* DH5α (Invitrogen catalog no. 18258-012) with heat shock method. Cells were incubated with plasmid library for 5 min on ice and were subjected to 42°C for 30 sec. Then cells were transferred back to ice and incubated for 2 min. Next, 0.95 mL SOC medium was added to the cells and incubated at 37°C for 1 h by shaking at 225 rpm. Cells were plated on LB-agar plates containing 100 µg/ml ampicilin. Plates were incubated for 16h at 37°C to grow the colonies and 3500 (approx 3-fold of library size) colonies were collected with liquid LB. Cells were transferred into 100 mL liquid LB and incubated at 37°C for 2 h. Plasmids were purified using Qiagen maxiperep kit (catalog no. 12163), according to the manufacturer’s protocol.

### In vitro transcription from plasmid library

Plasmids were linearized by NotI-HF enzyme so that the SP6 polymerase promoter site will be upstream of the sequences to be transcribed. Reactions consists of 5 U NotI-HF (NEB catalog no. R3189L), 5 µg library plasmid DNA, 1 X NEBuffer 4 (supplied with enzyme) and 90 µl of dH_2_O. Reaction was incubated at 37°C for 2 h to achieve complete digestion. The complete digestion of plasmid DNA was assessed by DNA gel electrophoresis. To eliminate NotI-HF and possible RNase in reaction mixture, samples was subjected to Proteinase K treatment. SDS and Proteinase K were added to the reaction mixture to a final concentration of 0.5% and 100 µg/mL, respectively. Sample was incubated at 37°C for 30 min. After Proteinase K treatment, sample was subjected to the phenol/chlororform extraction, followed by ethanol precipitation. Pellet was dissolved in 50 µl of RNase-free water. Next in vitro transcription was carried out using MAXIscript® SP6 Kit (Ambion catalog no: AM1308). Reaction composed of 1 µg of library plasmid, 1X transcription buffer, 0.5 mM of NTPs (GTP, ATP, CTP, and UTP), 40 U of SP6 RNA polymerase and 10 µl of RNase-free water. Reaction was incubated at 37°C for 30 min. Next, samples were treated with TURBO DNase to remove the plasmid templates. Briefly, 10 U of TURBO DNase (included with MAXIscript SP6 kit) were added to reaction mixture and incubated at 37°C for 15 min. To stop the reaction 1 μL of 0.5 M EDTA was added. To remove unincorporated NTPs and other impurities sample was precipitated with ammonium acetate/ethanol. The following reagents were added to the DNase-treated reaction mixture: 30 μL RNase-free water to bring the volume to 50 μL, 5 μL 5 M Ammonium Acetate, and 3 volumes 100% ethanol. Sample was chilled at −20°C for 30 min and then centrifuged at maximum speed in a 4°C table-top microfuge. The supernatant was discarded and pellet was washed with ice-cold, 70% ethanol. Pellet was dissolved in 50 μL RNase-free water and quality of RNA was assessed by agrose gel electrophoresis. In addition, PCR was carried out with in vitro transcribed RNA to confirm total depletion of plasmid DNA as well.

### Mouse liver collection and RNA extraction

WT 6-week old male C57/BL6 mice were acquired from Jackson Labs. Mice were sacrificed and liver samples were quickly dissected and snap-frozen in liquid nitrogen. RNA was isolated from frozen mouse liver samples by TRIzol reagent according to manufacturer’s protocol (Invitrogen catalog no. 15596-026). All animal experiments were performed in accordance with the approval of the Institutional Animal Care and Use Committee.

### Construction and sequencing of RNA-seq library from IVT RNA

IVT RNA (2500 ng, 150 ng, 75 ng, 15 ng, and 0 ng) was pooled with mouse liver RNA (0 ng, 2350 ng, 2425 ng, 2485 ng, and 2500 ng respectively) to a final quantity of 2.5 µg. Each pool was split into two replicate samples of 1 µg each. RNA pools were treated with Ribo-Zero Gold kit (Epicentre catalog no. RZHM11106) and converted into Illumina RNA-seq libraries with the TruSeq RNA sample prep kit (Ilumina catalog no. FC-122-1001). Briefly, rRNA was removed from 1 ug of pooled RNA using Ribo-Zero Gold kit and purified via ethanol/sodium acetate precipitation according to manufacturer’s protocol. After drying, the RNA pellet was dissolved in 18 μL of Elute, Prime, Fragment mix (provided with TruSeq RNA sample prep kit). RNA was fragmented for 8 minutes and 17 uL of this fragmented RNA was used to make the RNA-seq library according to Illumina TruSeq RNA sample prep kit protocol. After fragmentation/ priming, first strand cDNA synthesis with SuperScript II (Invitrogen catalog no. 18064014), second-strand synthesis, end-repair, a-tailing, and adapter ligation, the library fragments were enriched with 15 cycles of PCR. Quality and size of library was assessed using Agilent 2100 BioAnalyzer. The five libraries from each replicate were pooled together and sequenced using a single lane from an Illumina HiSeq 2000 (paired 100 bp reads).

### Construction and sequencing of plasmid library

MGC plasmids were linearized by NotI-HF enzyme as before. These linearized plasmids were then fragmented using a Covaris S220 Focused-ultrasonicator. Briefly, 1.2 µg of linearized plasmid in a final volume of 60 uL of H_2_O was loaded into a microTUBE (Covaris catalog no. 520045). The ultrasonicator was de-gassed and prepared according to manufacturer’s protocol. Linearized plasmids were sonicated using the following conditions: intensity 5, duty factor 10%, cycles per burst 200, time 120s, and water bath temperature 7°C. Fragmented plasmids were gel-purified using a 1% agarose gel (BioRad catalog no. 161-3107) and TAE running buffer (BioRad catalog no. 161-0743). Gel slice between 100 bp and 700 bp was excised and DNA was purified using MinElute gel extraction kit (Qiagen catalog no. 28606) according to manufacturer’s protocol. Fragmented DNA was converted into a sequencing library using TruSeq DNA sample prep kit (Illumina catalog no. FC-121-2001). End repair, adenylation, adapter ligation, gel size-selection, and PCR enrichment were performed according to manufacturer’s protocol. During the gel size-selection, a band between 300 bp and 500 bp was excised. Quality and size of library was assessed using Agilent 2100 BioAnalyzer. This library was sequenced using an Illumina MiSeq (paired 100 bp reads).

### Construction and sequencing of no selection and polyA libraries

As with the other RNA-seq libraries, these libraries were prepared using the TruSeq RNA sample prep kit (Illumina catalog no. FC-122-1001). For the polyA sample, 1 µg of IVT RNA was treated with polyA selection reagents included with the TruSeq RNA sample prep kit according to manufacturer’s protocol. The remainder of the library preparation was carried out using the same conditions as for the other IVT RNA samples. For the no selection sample, 100 ng of IVT RNA at a concentration of 100 ng/µL was diluted with 17 μL of Elute, Prime, Fragment mix (provided with TruSeq RNA sample prep kit). Again, the remainder of the library preparation was carried out as with the other samples. These samples were sequence in a single Illumina HiSeq 2500 lane (paired 100 bp reads).

### Aligning, quantifying, and visualizing sequencing data

Raw reads from all sequencing samples were aligned to the human genome (GRCh37/hg19) using the RNA-seq Unified Mapper [14] (RUM; v2.0.4) with default parameters. RUM also generated RNA-seq coverage plots in bedgraph format, and calculated transcript- and exon-level FPKM values for each IVT transcript (accession numbers listed in Additional file 10). All analyses were performed using uniquely aligned reads (no multi-mappers) from the RUM_Unique and RUM_Unique.cov output files. Quantification was performed using annotations for the IVT transcripts that we downloaded from the MGC Genes track [17] on the UCSC Genome Browser [23]. Those IVT transcripts mapping to multiple loci, or overlapping other IVT transcripts were removed from further analysis (marked with * in Additional file 10). All coverage plots in this paper were visualized in and captured from the UCSC Genome Browser. All statistical tests and correlation plots were performed in R.

### Generating simulated data

Simulated data was generated using the BEERS software package (http://www.cbil.upenn.edu/BEERS/) from gene models for IVT transcripts, with an average coverage depth of 1000 reads. All error, intronic read, and polymorphism parameters were set to zero. Remaining parameters used default values. For the QM-simulated (Quantity Matched) data, FPKM values from replicate 1 of the IVT-only data were used as seeds for generating expression levels. All other parameters were the same as for the other simulated data.

### Processivity analysis

Coverage data for each IVT transcript was extracted from coverage files for the plasmid and no selection samples. For each transcript, base pair-level coverage data was extracted from the regions spanning 5-15% and 85-95% of the transcript, by length. For example, given a 1000 bp transcript, the first region spans base pairs 50-150, and the second region spans base pairs 850-950. These two coverage regions represent the 5’ end and 3’ end of the transcript, respectively. The first and last 5% of the transcript was excluded to avoid artifacts from the fragmentation process. Processivity of each transcript was assessed by the ratio of the mean depth of coverage from both of these regions (3’ region mean / 5’ region mean). These processivity ratios were calculated for all transcripts in the plasmid and no selection data, with expression > 5FPKM.

### Calculating fold-change difference in within-transcript coverage

Coverage data for each of the IVT transcripts was extracted from the coverage files for the IVT-only, polyA, and no selection samples. The first and last 200 bp were trimmed from each transcript to prevent edge effects from interfering with the calculations. Due to this trimming, all IVT transcripts with less than 500 bp were discarded. Nucleotide-level coverage data was grouped into percentiles based on depth of coverage. Average coverage across the 10^th^ percentile and 90^th^ percentile were calculated. Fold-change difference in within-transcript coverage were calculated by dividing the 90^th^ percentile average by the 10^th^ percentile average. The list of transcripts with associated fold-change values is included in Additional file 13.

### Identifying hunc regions

Coverage data for each of the IVT transcripts was extracted from the coverage files from each of the rRNA-depleted datasets (replicate dilution series: IVT-only, 1 IVT: 1 mouse, 1 IVT: 2 mouse, 1 IVT: 10 mouse, and mouse-only). These coverage plots were normalized between 0 and 1 to allow comparison between different dilutions. For each nucleotide position in a transcript, the deviation in coverage between each of the samples was calculated using the median absolute deviation (MAD), due to its resistance to outliers. MAD scores were calculated across the different dilutions using R’s *mad* function with constant = 1. Next, a sliding window was used to calculate the average MAD in the 100 bp windows centered on each nucleotide in the transcript. The first 300 and last 250 windows were trimmed from each transcript to avoid confounding variability due to edge effects or fragmentation artifacts. All analysis up until this point was carried out separately on the two replicate datasets. The 95^th^ percentile of MAD scores was calculated for each of the replicates using R’s *quantile* function (replicate 1: 0.08810424, replicate 2: 0.07183765). Only those regions with at least 20 contiguous windows having MAD scores above the appropriate 95^th^ percentile values were retained for further analysis. Next, the BEDTools [24] intersect function was used to remove any regions with high MAD scores not present in both replicates. Finally, these remaining regions of high coverage variability were filtered for mouse reads misaligned to the human reference. Any region coming from a transcript with FPKM >= 5 in the mouse-only samples were discarded. To account for localized misalignment of mouse reads, any regions with an average coverage > 10 in the mouse-only samples or in the 100 bp on either side of the region were discarded. These remaining regions comprise the list final list of regions with high coverage variability. To search for hunc regions not maintained between replicates, windowed MAD scores from replicate 2 were subtracted from those of replicate 1. The 2.5^th^ and 97.5^th^ percentiles of these difference values were used as cutoffs (2.5^th^ percentile: −0.07053690, 97.5^th^ percentile: 0.09134876) to pull out the most extreme positive and negative difference values. Regions corresponding to these extreme difference values were filtered as above. Additionally, those difference regions within 200 bp of a previously identified hunc regions were filtered out. This last filtering step accounts for cases where a difference region with high MAD scores is just an extension of an existing hunc region. Hunc regions and difference regions were manually checked to determine whether or not they represent regions where expression patterns deviate from the remainder of the transcript.

### Generating sequence characteristics

Sequences for each transcript were collected in R using the BSgenome, GenomicRanges, and GenomicFeatures packages. Hexamer entropy for each transcript was calculated as follows: occurrences of all possible hexamers in a given transcript were counted. These counts were converted into frequency space, and these frequency values were used to calculate the Shannon entropy. Shannon entropy is commonly used to represent complexity in nucleotide sequences or multiple alignments [19–21]. Similarity of transcripts to rRNA sequences were calculated as follows: each transcript was aligned to 45S (NR_046235.1) and 5S (X71804.1) rRNA using NCBI-BLAST [25] and the e-score for the best alignment was saved.

### Sequence characteristic analysis

The list of IVT transcripts were sorted by transcript-level coefficients of variation for each library method (plasmid, no selection, polyA, replicate 1 of rRNA-depleted IVT-only). All transcripts with transcript-level FPKM <= 5 were excluded from further analysis. From this sorted list the transcripts with the 100 least and 100 most extreme coefficients of variation were collected for each of the above sequencing samples. The values for hexamer entropy, GC-content, and rRNA sequence similarity were compared between every pair of 100 least and 100 most extreme coefficients of variation using a Wilcoxon signed-rank test (implemented in R as the wilcox.test function). This entire analysis was repeated using transcript-level FPKM values instead of the coefficients of variation. All boxplots were prepared using R.

## Data access

We deposited all sequencing data in the NCBI Gene Expression Omnibus (GEO) under accession number GSE50445. Reviewer data link: http://www.ncbi.nlm.nih.gov/geo/query/acc.cgi?token=zdyfjmiagoacglq&acc=GSE50445

## Acknowledgements

We would like to thank the Penn Genome Frontiers Institute sequencing core, the Institute for Diabetes, Obesity and Metabolism, the DRC grant (P30DK19525), and the services of the Functional Genomics Core for performing the Illumina sequencing. J.B.H. is supported by the National Institutes of Health grants 2-R01-NS054794-06 and 5-R01-HL097800-04 and by DARPA [12-DARPA-1068] (to John Harer, Duke University). G.R.G is supported by the National Center for Research Resources and the National Center for Advancing Translational Sciences, National Institutes of Health, through Grant UL1TR000003. This project is funded, in part, by the Penn Genome Frontiers Institute under a HRFF grant with the Pennsylvania Department of Health, which disclaims responsibility for any analyses, interpretations or conclusions. This project is also supported in part by the Institute for Translational Medicine and Therapeutics of the Perelman School of Medicine at the University of Pennsylvania. He content is solely the responsibility of the authors and does not necessarily represent the official views of the NIH.

## Authors’ contributions

J.B.H. and R.I. conceived the research. N.F.L. and I.H.K. created the Illumina sequencing libraries. N.F.L., G.R.G., M.B., R.T., and R.Z., developed and performed the computational analysis. H.D. and A.P. contributed code. R.I., J.K., H.D., K.H., and A.P assisted with the computational analysis. J.B.H., G.R.G., and N.F.L. wrote the paper.

## Additional data files

### Additional_file_1 as PDF

**Additional file 1: Figure S1.** Expression comparison between replicates. (A) Correlation plots for log10 transcript-level FPKM values between replicate IVT-seq samples. Pearson R^2^ values for the correlations are included as inserts in each plot. (B) Distribution of FPKM values in both replicates of the IVT-only sample. FPKM values are plotted on the x-axis in log10 space. The y-axis is plotted in arbitrary density units.

### Additional_file_2 as PDF

**Additional file 2: Figure S2.** Expression comparison between simulated and IVT data. Correlation plots for log10 transcript-level FPKM values between (A) simulated data or (B) QM-simulated data, and replicate 1 of the IVT-only data. Pearson R^2^ values for the correlations are included as inserts in each plot.

### Additional_file_3 as PDF

**Additional file 3: Figure S3.** Coverage patterns are reproducible across replicates. Coverage patterns from both replicates for all transcripts in Figure 2. RNA-seq coverage plots from replicate IVT only samples (red – replicate 1l blue – replicate 2) for (A) BC015891, (B) BC009037, and (C) BC016283 are displayed according to the gene model (green), as it is mapped to the human reference genome. Blocks correspond to exons and lines indicate introns. The chevrons within the intronic lines indicated the direction of transcription. Numbers on y-axis refer to RNA-seq read-depth at a given nucleotide position. All transcripts are displayed in the 5′ to 3′ direction.

### Additional_file_4 as PDF

**Additional file 4: Figure S4.** Fold-change in within-transcript coverage across libraries. The cumulative distribution functions for fold-change in within transcript coverage are displayed for the rRNA-depleted (red), polyA (orange), no selection (green), plasmid (blue), QM-simulated (gray), and simulated (black) datasets. Curves toward the left side of the plot indicate fewer genes contain high fold-change differences in coverage. Curves toward the right side of the plot indicate many genes contain high fold-change differences in coverage. The dotted lines indicate the y-axis values for none of the data (0.0) and all of the data (1.0). This plot is focused on the fold-change values between 1 and 10. See the *Materials and Methods* section for full details on the fold-change calculations.

### Additional_file_5 as PDF

**Additional file 5: Figure S5.** Plasmid sequencing protocol compared to IVT-seq. The protocol for preparing MGC plasmids for DNA-sequencing library generation is displayed alongside the protocol for preparing IVT transcripts for RNA-seq library generation. Both protocols start by linearizing the plasmids. For DNA-sequencing, linearized plasmids are fragmented via Covaris sonication, and the resulting fragments are taken through the TruSeq protocol. For RNA-sequencing, the linearized plasmids are used as templates for an *in vitro* transcription reaction. IVT RNA is then pooled with mouse RNA, rRNA is removed from pool via Ribo-Zero Gold kit, rRNA-depleted pool is fragmented via metal-ion hydrolysis, and fragmented RNA is converted to cDNA via reverse transcription with random-hexamer priming. The resulting cDNA fragments are then taken through the TruSeq protocol.

### Additional_file_6 as PDF

**Additional file 6: Figure S6.** Nucleotide frequency as a function of read position for sequencing reads at the 5’ ends of cDNA fragments. Frequencies are plotted for plasmid, no selection, rRNA-depleted, and polyA datasets.

### Additional_file_7 as PDF

**Additional file 7: Figure S7.** Confounding effects between coverage depth and variability. Distributions of transcript-level coefficients of variation for the 100 transcripts with the highest and lowest transcript-level FPKMs from the plasmid, no selection, rRNA-depleted, and polyA libraries. Asterisks indicate the significance of a Wilcoxon signed-rank test comparing values for the listed sequence characteristics between each pair of groups from the same sample. * = *p*-value < 0.05; *** = *p*-value < 0.001.

### Additional_file_8 as PDF

**Additional file 8: Figure S8.** Effects of sequence characteristics on coverage depth. Distributions of (A) hexamer entropy, (B) GC-content, and (C) rRNA sequence similarity for the 100 transcripts with the highest and lowest transcript-level FPKMs from the plasmid, no selection, rRNA-depleted, and polyA libraries. Asterisks indicate the significance of a Wilcoxon signed-rank test comparing values for the listed sequence characteristics between each pair of groups from the same sample. ** = *p*-value < 0.01; *** = *p*-value < 0.001.

### Additional_file_9 as PDF

**Additional file 9: Figure S9.** rRNA sequence similarity and coverage bias in rRNA-depleted data. Correlation plot between Smith-Waterman alignment score to rRNA sequences and the magnitude of the decrease in coverage depth between no selection and rRNA-depleted samples. A coverage drop of 1.0 indicates a large decrease in coverage between the no selection and rRNA-depleted samples. A coverage drop of 0 indicates no difference between the two samples. For full details on this analysis, see Additional file 12.

### Additional_file_10 as XLSX

Additional file 10: Accession numbers for IVT transcripts.

### Additional_file_11 as BED

Additional file 11: List of regions with high coverage variability.

### Additional_file_12 as PDF

Additional file 12: Description of window analysis of rRNA sequence similarity.

### Additional_file_13 as XLSX

Additional file 13: List of transcripts with associated fold-change values in within-transcript coverage.

## References

1. Wang Z, Gerstein M, Snyder M: RNA-Seq: a revolutionary tool for transcriptomics. Nat Rev Genet 2009, 10:57–63.

2. Park E, Williams B, Wold BJ, Mortazavi A: RNA editing in the human ENCODE RNA-seq data. Genome Res 2012, 22:1626–1633.

3. Ilott NE, Ponting CP: Predicting long non-coding RNAs using RNA sequencing. Methods San Diego Calif 2013.

4. Nagalakshmi U, Waern K, Snyder M: RNA-Seq: a method for comprehensive transcriptome analysis. Curr Protoc Mol Biol Ed Frederick M Ausubel Al 2010, Chapter 4:Unit 4.11.1–13.

5. Levin JZ, Yassour M, Adiconis X, Nusbaum C, Thompson DA, Friedman N, Gnirke A, Regev A: Comprehensive comparative analysis of strand-specific RNA sequencing methods. Nat Methods 2010, 7:709–715.

6. Cui P, Lin Q, Ding F, Xin C, Gong W, Zhang L, Geng J, Zhang B, Yu X, Yang J, Hu S, Yu J: A comparison between ribo-minus RNA-sequencing and polyA-selected RNA-sequencing. Genomics 2010, 96:259–265.

7. Adiconis X, Borges-Rivera D, Satija R, DeLuca DS, Busby MA, Berlin AM, Sivachenko A, Thompson DA, Wysoker A, Fennell T, Gnirke A, Pochet N, Regev A, Levin JZ: Comparative analysis of RNA sequencing methods for degraded or low-input samples. Nat Methods 2013, 10:623–629.

8. Benjamini Y, Speed TP: Summarizing and correcting the GC content bias in high-throughput sequencing. Nucleic Acids Res 2012.

9. Aird D, Ross MG, Chen W-S, Danielsson M, Fennell T, Russ C, Jaffe DB, Nusbaum C, Gnirke A: Analyzing and minimizing PCR amplification bias in Illumina sequencing libraries. Genome Biol 2011, 12:R18.

10. Hansen KD, Brenner SE, Dudoit S: Biases in Illumina transcriptome sequencing caused by random hexamer priming. Nucleic Acids Res 2010, 38:e131–e131.

11. Nakamura K, Oshima T, Morimoto T, Ikeda S, Yoshikawa H, Shiwa Y, Ishikawa S, Linak MC, Hirai A, Takahashi H, Altaf-Ul-Amin M, Ogasawara N, Kanaya S: Sequence-specific error profile of Illumina sequencers. Nucleic Acids Res 2011.

12. Spicuglia S, Maqbool MA, Puthier D, Andrau J-C: An update on recent methods applied for deciphering the diversity of the noncoding RNA genome structure and function. Methods.

13. Trapnell C, Williams BA, Pertea G, Mortazavi A, Kwan G, van Baren MJ, Salzberg SL, Wold BJ, Pachter L: Transcript assembly and quantification by RNA-Seq reveals unannotated transcripts and isoform switching during cell differentiation. Nat Biotechnol 2010, 28:511–515.

14. Grant GR, Farkas MH, Pizarro AD, Lahens NF, Schug J, Brunk BP, Stoeckert CJ, Hogenesch JB, Pierce EA: Comparative analysis of RNA-Seq alignment algorithms and the RNA-Seq unified mapper (RUM). Bioinforma Oxf Engl 2011, 27:2518–2528.

15. Lee S, Seo CH, Lim B, Yang JO, Oh J, Kim M, Lee S, Lee B, Kang C, Lee S: Accurate quantification of transcriptome from RNA-Seq data by effective length normalization. Nucleic Acids Res 2011, 39:e9–e9.

16. Jiang L, Schlesinger F, Davis CA, Zhang Y, Li R, Salit M, Gingeras TR, Oliver B: Synthetic spike-in standards for RNA-seq experiments. Genome Res 2011, 21:1543–1551.

17. Temple G, Gerhard DS, Rasooly R, Feingold EA, Good PJ, Robinson C, Mandich A, Derge JG, Lewis J, Shoaf D, Collins FS, Jang W, Wagner L, Shenmen CM, Misquitta L, Schaefer CF, Buetow KH, Bonner TI, Yankie L, Ward M, Phan L, Astashyn A, Brown G, Farrell C, Hart J, Landrum M, Maidak BL, Murphy M, Murphy T, Rajput B, et al.: The completion of the Mammalian Gene Collection (MGC). Genome Res 2009, 19:2324–2333.

18. Bailey TL, Boden M, Buske FA, Frith M, Grant CE, Clementi L, Ren J, Li WW, Noble WS: MEME SUITE: tools for motif discovery and searching. Nucleic Acids Res 2009, 37(Web Server issue):W202–208.

19. Schmitt AO, Herzel H: Estimating the entropy of DNA sequences. J Theor Biol 1997, 188:369– 377.

20. Mantegna RN, Buldyrev SV, Goldberger AL, Havlin S, Peng CK, Simons M, Stanley HE: Linguistic features of noncoding DNA sequences. Phys Rev Lett 1994, 73:3169–3172.

21. Koonin EV: A non-adaptationist perspective on evolution of genomic complexity or the continued dethroning of man. Cell Cycle Georget Tex 2004, 3:280–285.

22. Dohm JC, Lottaz C, Borodina T, Himmelbauer H: Substantial biases in ultra-short read data sets from high-throughput DNA sequencing. Nucleic Acids Res 2008, 36:e105.

23. Rhead B, Karolchik D, Kuhn RM, Hinrichs AS, Zweig AS, Fujita PA, Diekhans M, Smith KE, Rosenbloom KR, Raney BJ, Pohl A, Pheasant M, Meyer LR, Learned K, Hsu F, Hillman-Jackson J, Harte RA, Giardine B, Dreszer TR, Clawson H, Barber GP, Haussler D, Kent WJ: The UCSC Genome Browser database: update 2010. Nucleic Acids Res 2010, 38(Database issue):D613–619.

24. Quinlan AR, Hall IM: BEDTools: a flexible suite of utilities for comparing genomic features. Bioinforma Oxf Engl 2010, 26:841–842.

25. Altschul SF, Gish W, Miller W, Myers EW, Lipman DJ: Basic local alignment search tool. J Mol Biol 1990, 215:403–410.

